# Signaling to TRP53 and TAp63 from CHK1/CHK2 is responsible for elimination of most oocytes defective for either chromosome synapsis or recombination

**DOI:** 10.1101/768150

**Authors:** Vera D. Rinaldi, Jordana C. Bloom, John C. Schimenti

**Author notes:** These authors contributed equally to the manuscript (Vera D. Rinaldi, Jordana C. Bloom). Corresponding author (J.C.S.).

## Abstract

Eukaryotic organisms have evolved mechanisms to prevent the accumulation of cells bearing genetic aberrations. This is especially crucial for the germline, because fecundity, and fitness of progeny would be adversely affected by an excessively high mutational incidence. The process of meiosis poses unique problems for mutation avoidance, due to the requirement for SPO11-induced programmed double strand breaks (DSBs) in recombination-driven pairing and segregation of homologous chromosomes. Mouse meiocytes bearing unrepaired meiotic DSBs or unsynapsed chromosomes are eliminated before completing meiotic prophase I. In previous work, we showed that checkpoint kinase 2 (CHK2; CHEK2), a canonical DNA damage response protein, is crucial for eliminating not only oocytes defective in meiotic DSB repair (e.g. *Trip13*^*Gt*^ mutants), but also asynaptic *Spo11*^−/−^ oocytes that accumulate a threshold level of spontaneous DSBs. However, rescue of such oocytes by *Chk2* deficiency was incomplete, raising the possibility that a parallel checkpoint pathway(s) exists. Here, we show that mouse oocytes lacking both TAp63 and TRP53 protects nearly all *Spo11*^−/−^ and *Trip13*^*Gt/Gt*^ oocytes from elimination. We present evidence that checkpoint kinase I (CHK1; CHEK1), which is known to signal to TRP53, also becomes activated by persistent DSBs in oocytes, and to an increased degree when CHK2 is absent. The combined data indicate that nearly all oocytes reaching a threshold level of unrepaired DSBs are eliminated by a semi-redundant pathway of CHK1/CHK2 signaling to TRP53/TAp63.

## INTRODUCTION

Meiocytes from diverse organisms have developed mechanisms for minimizing the production of gametes that containing genetic anomalies such as unrepaired DSBs and homologous chromosome asynapsis. Mouse oocytes bearing mutations that prevent repair of programmed SPO11/TOPOVIBL-induced DSBs, which are essential for recombination-mediated pairing and synapsis of homologous chromosomes (Baudat *et al.* 2000; Romanienko and Camerini-Otero 2000; Mahadevaiah *et al.* 2001; Robert *et al.* 2016), are eliminated by a DNA damage checkpoint (Di Giacomo *et al.* 2005). The molecular nature of this checkpoint was first revealed as involving signaling of CHK2 to TRP53 and TP63 (the TA isoform, denoted here as TAp63), by studies exploiting a hypomorphic allele (*Trip13*^*Gt*^) of *Trip13* that causes sterility (Bolcun-Filas *et al.* 2014). This allele was useful because it is defective for DSB repair but not synapsis (Li and Schimenti 2007). Deficiency of *Chk2* protected against oocyte loss and restored fertility of *Trip13*^*Gt/Gt*^ females. *Chk2* also plays a role in the DNA damage checkpoint in spermatocyte meiosis (Pacheco *et al.* 2015).

Defects in chromosome synapsis during meiotic prophase I also triggers death of most oocytes. There are at least two mechanisms underlying this “synapsis checkpoint.” One is meiotic silencing of unsynapsed chromatin (MSUC), a process of extensive heterochromatinization and transcriptional downregulation, which appears to function primarily in situations where only ~1-3 chromosomes are asynapsed (Kouznetsova *et al.* 2009; Cloutier *et al.* 2015). A second mechanism pertains to oocytes that are highly asynaptic, in which the silencing machinery is presumably overwhelmed (Kouznetsova *et al.* 2009). Surprisingly, this mechanism is also highly dependent on the DNA damage checkpoint. The mechanistic basis for this is the formation of a threshold level (~10) of spontaneously-arising DSBs that are SPO11-independent (Carofiglio *et al.* 2013; Rinaldi *et al.* 2017). Approximately 61% of *Spo11*^−/−^ oocytes, which lack any appreciable degree of homolog synapsis and are incapable of forming programmed meiotic DSBs, reach this threshold, and almost the entire ovarian reserve (primordial oocytes) is depleted by 3 weeks after birth. *Chk2* deletion rescued oocyte numbers to ~25% of WT, indicating that most are either eliminated by an alternative pathway, or that they succumb nonspecifically from a catastrophically high number of DSBs (up to ~100, with an average of ~50/cell) (Rinaldi *et al.* 2017). Similarly, *Chk2* deficiency rescued *Trip13*^*Gt/Gt*^ oocytes to ~1/3 of WT levels (Bolcun-Filas *et al.* 2014), raising the possibility that the same CHK2-independent pathway may be active in both cases. Here, we tested the possibility that the cause for incomplete rescue is the existence of another pathway that is distinct or complementary to that involving CHK2, but which also involves signaling to TRP53 and TAp63. Our results indicate that this is indeed the case, and that most *Spo11*^−/−^ and TRIP13-deficient oocytes are ultimately eliminated by the combined activation of TRP53 and TAp63.

## RESULTS and DISCUSSION

To address the question of whether a CHK2-independent pathway exists that can eliminate oocytes bearing unrepaired DSBs, we utilized two mutant models, *Trip13*^*Gt*^ and a *Spo11* null (*Spo11*^−^). Virtually all *Trip13*^*Gt/Gt*^ oocytes are eliminated due to failure to repair SPO11-dependent DSBs (Li and Schimenti 2007) by the end of pachynema (Rinaldi *et al.* 2017). *Chk2* deficiency rescued ~ 1/3 of these oocytes, and these rescued oocytes gave rise to viable offspring (Bolcun-Filas *et al.* 2014). Although nullizygosity for either of CHK2’s downstream phosphorylation targets, *Trp53* and *TAp63*, enabled little or no rescue of *Trip13*^*Gt/Gt*^ oocytes, *Trip13*^*Gt/Gt*^ *TAp63*^−/−^ *Trp53*^+/−^ mice exhibited oocyte rescue to a degree similar to that of *Trip13*^*Gt/Gt*^ *Chk2*^−/−^ mice (Bolcun-Filas *et al.* 2014). At the time of that report, double nulls (*TAp63*^−/−^ *Trp53*^−/−^) were not assayed for the extent to which they could rescue *Trip13*^*Gt/Gt*^ oocytes. We hypothesized that the inability to achieve full oocyte rescue in either *Trip13*^Gt/Gt^ TAp63^−/−^ *Trp53*^+/−^ or *Trip13*^*Gt/Gt*^ *Chk2*^−/−^ females was due to one of the following: 1) the number of DSBs was so high in most oocytes that elimination occurred in a checkpoint-independent fashion; 2) residual TRP53 activity in the *Trip13*^*Gt/Gt*^ *TAp63*^−/−^ *Trp53*^+/−^ mice sufficed to trigger apoptosis in many oocytes; and/or 3) a parallel checkpoint pathway is active in *Chk2*^−/−^ oocytes.

To test these possibilities, we first bred and assessed the ovarian reserve in *Trip13*^*Gt/Gt*^ *Trp53*^−/−^ *TAp63*^−/−^ mice. Remarkably, the numbers of primordial and more developed oocytes in the triple mutants were indistinguishable from WT (Fig. 1a,b). This result indicates that essentially all *Trip13*^*Gt/Gt*^ oocytes are eliminated by checkpoint signaling to TRP53 and TAp63, thereby eliminating hypothesis 1, but supporting hypothesis 2. This result is also consistent with hypothesis 3, implying that another pathway or kinase is signaling to these two effector proteins.

**Figure 1.**
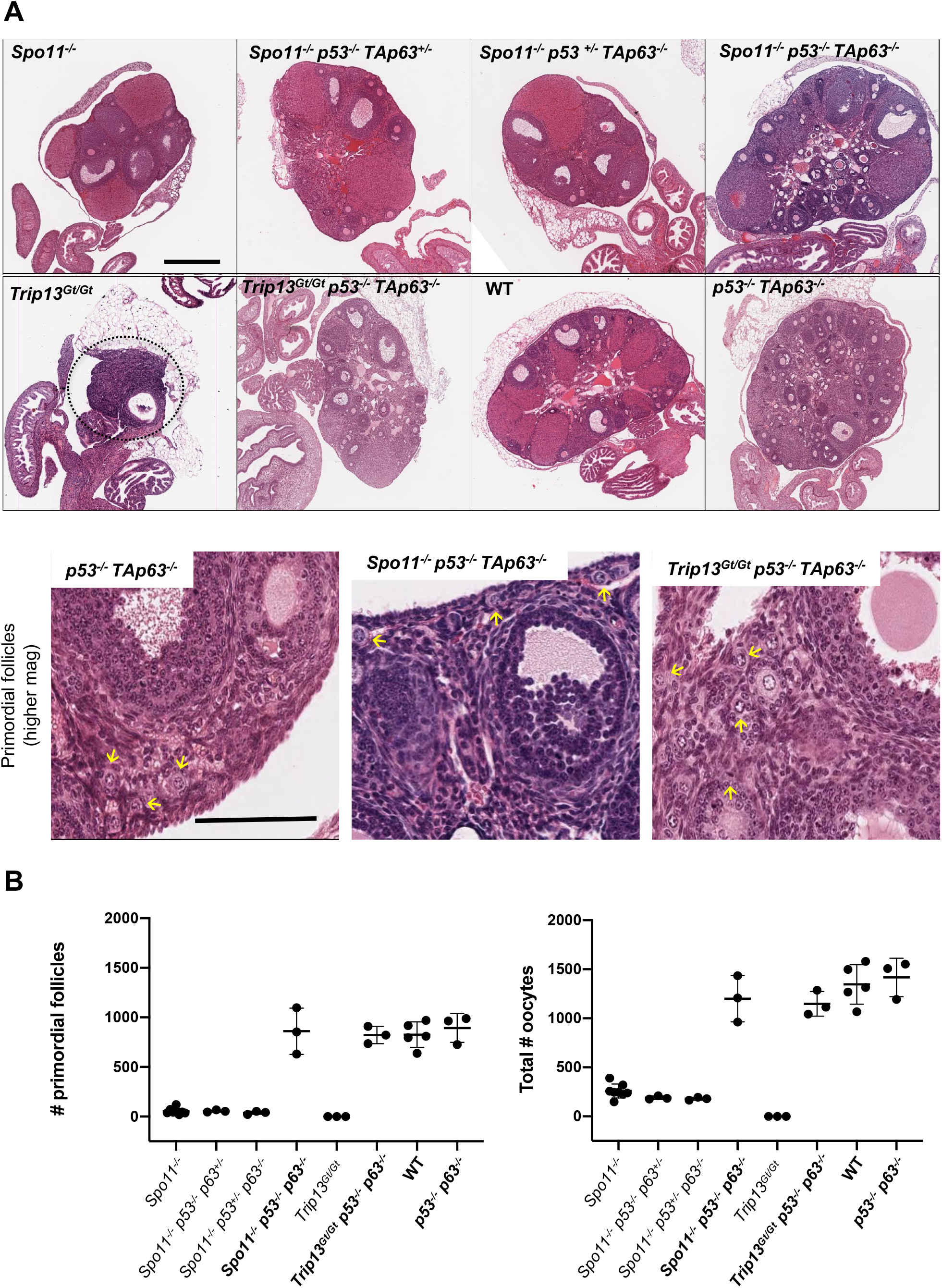
Rescue of SPO11- and TRIP13-deficient oocytes by compound deletion of *p53* and *TAp63*. (A) Hematoxylin and eosin stained ovaries from 2-month-old mice of the indicated genotypes. The top two rows are images are from histological sections through the approximate center of the ovaries. Scale bar = 500 µM. The dashed circle indicates the residual *Trip13*^*Gt/Gt*^ ovary. The bottom row shows higher magnification (scale bar = 100µM) images of selected genotypes. Yellow arrows indicate examples of primordial follicles. (B) Oocyte quantification. To the left is the quantification of primordial follicles (p<0.0001 for all oocyte rescued (in bold) genotypes compared to non-rescued genotypes; p=0.92 for oocyte rescued genotypes vs. WT and *Trp53*^−/−^ *TAp63*^−/−^ ovaries) and total oocytes (from all stages of follicles) from individual ovaries (p<0.0001 for all oocyte rescued genotypes compared to non-rescued genotypes; p=0.33 for oocyte rescued genotypes vs. WT and *Trp53*^−/−^ *TAp63*^−/−^ ovaries). Genotype abbreviations are as follows: *TAp63* is abbreviated as *p63*; WT = wild-type.

Next, we tested whether the incomplete rescue of *Spo11*^−/−^ oocytes by *Chk2* deletion is also potentially a consequence of checkpoint signaling to TAp63 and TRP53 via a different transducer. Accordingly, we bred mice that lacked either or both of these proteins in the context of *Spo11* deficiency. Oocyte numbers in *Spo11*^−/−^ mice that were also homozygous for mutations in either *Trp53* or *TAp63* and heterozygous for a mutation in the other (*Trp53*^−/−^ *TAp63*^+/−^ and *Trp53*^+/−^ *TAp63*^−/−^) were indistinguishable from *Spo11* nulls; nearly the entire oocyte reserve was depleted after 2 months of age, as is characteristic for *Spo11* deficiency (Di Giacomo *et al.* 2005). However, homozygosity for both *Trp53* and *TAp63* dramatically restored oocyte numbers to WT levels (Fig. 1a,b).

These experiments indicate that unrepaired meiotic DSBs, when present at levels above the threshold to trigger their elimination (Rinaldi *et al.* 2017), ultimately cause DNA damage signaling to TRP53 and TAp63. Additionally, we conclude that one or both of these proteins can be activated not only by CHK2, but also another kinase. In our previous studies, we suggested that the apical kinase ATM, which when activated by DSBs typically phosphorylates CHK2, is not essential for the meiotic DNA damage checkpoint (Bolcun-Filas *et al.* 2014). This conclusion was based on the observation that many *Atm*^−/−^ oocytes, which have extensive DSBs due to the role of ATM in negatively regulating SPO11 (Lange *et al.* 2011), are eliminated in a CHK2-dependent manner. We proposed (Bolcun-Filas *et al.* 2014) that the related kinase ATR (ataxia telangiectasia and Rad3 related) might activate CHK2 in oocytes as has been shown in irradiated mitotic cells (Wang *et al.* 2006), which in turn would phosphorylate TAp63 and TRP53. Since ATR primarily activates CHK1, albeit most notably in the context of damage at DNA replication forks, we speculated that CHK1 can trigger death of DSB-bearing oocytes by activating TRP53 in the absence of CHK2. TRP53 is a known target of CHK1 (Shieh *et al.* 2000; Ou *et al.* 2005), and CHK1 can be activated in response to DSBs either in an ATM-dependent (Flaggs *et al.* 1997; Maréchal and Zou 2013) or – independent (Flaggs *et al.* 1997; Balmus *et al.* 2012) manner. Recombinant CHK1 was also reported to phosphorylate TRP63 *in vitro* (Kim *et al.* 2007).

If this hypothesis is true, CHK1 would be activated in response to DSBs present in oocytes. To test this, we examined levels of CHK1 phosphorylated at Ser345 (pCHK1; indicative of the active form) and TRP53 (which is stabilized by phosphorylation) in various genotypes of neonatal (3-5 days post-partum) ovaries treated with zero or 3Gy of ionizing radiation (IR). This level of IR induces ~40 DSBs, as measured by RAD51 foci (a proxy for DSBs) on oocyte meiotic chromosomes (Rinaldi *et al.* 2017). Three hours later, proteins were extracted for Western blot analysis. Because ovaries contain somatic cells, the germ cell marker MVH serves as a loading control. In unirradiated ovaries, there was no apparent difference between genotypes (WT; *Chk2*^−/−^; *Spo11*^−/−^; *Spo11*^−/−^ *Chk2*^−/−^) in the levels of pCHK1 or TRP53. Whereas exposure to IR caused a small increase of CHK1 phosphorylation in WT oocytes, the induction was substantial in oocytes deficient for CHK2 (Fig. 2; Fig. S1). This implies that the ATM and/or ATR kinases have a higher propensity to activate CHK2 than CHK1 in response to DSBs, but that the CHK1 pool becomes increasingly targeted in the absence of CHK2 until signaling downstream to TRP53/TAp63 is sufficient to trigger apoptosis or the damage is repaired.

**Figure 2.**
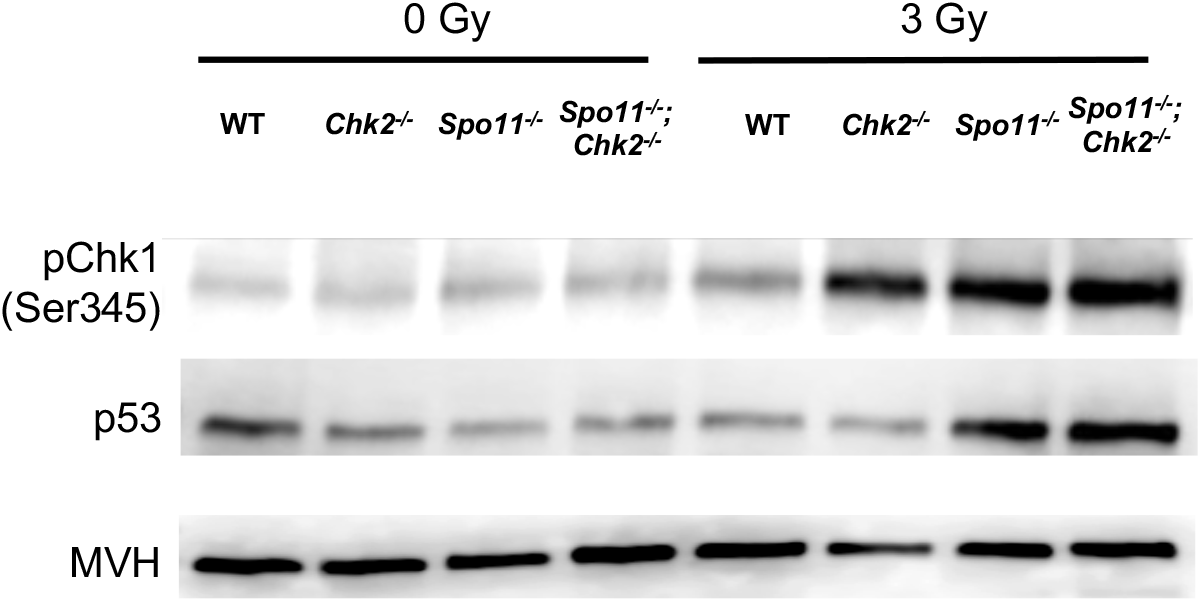
Increased CHK1 activation and p53 stabilization in CHK2-deficient oocytes. Western blot probed with indicated antibodies. Each lane contains total protein extracted from four ovaries (postnatal day 3-5) that were either exposed or not to ionizing radiation (IR). The same blot was stripped and re-probed sequentially. A biological replicate is shown in Fig. S1.

Interestingly, IR also caused a marked increase of pCHK1, compared to WT, in the in *Spo11*^−/−^ oocytes (Fig. 2; Fig. S1). Similarly, levels of p53 were also higher in IR-treated SPO11-deficient ovaries, but the presence or absence of CHK2 had no consequence (Fig. 2a). One possible explanation is that repair of IR-induced DSBs by intersister (IS) recombination is inhibited in *Spo11* mutants, because unsynapsed chromosome axes (synapsis is virtually absent in *Spo11*^−/−^ meiocytes) retain HORMAD1/2 proteins that prevent such repair (Carofiglio *et al.* 2013; Rinaldi *et al.* 2017). In contrast, *Chk2*^−/−^ oocytes would retain IS repair ability, and thus either delay or minimize signaling to TRP53. A second possible cause of increased pCHK1 in irradiated *Spo11*^−/−^ ovaries is that asynapsed chromosomes are more susceptible to IR-induced DNA damage than synapsed chromosomes (as in WT and *Chk2*^−/−^ oocytes). A final possibility is that the presence of ATR on asynapsed chromatin (Turner *et al.* 2004, 2006)(Perera *et al.* 2004; Cloutier *et al.* 2016) facilitates DNA damage signaling to CHK1 under conditions of unrepaired DSBs, implying that ATR is not only involved in MSUC, but also retains its function as key component of the DSB repair machinery (Widger *et al.* 2018).

In summary, we have shown that mouse oocytes with unrepaired DSBs or extensive asynapsis are culled by a DNA damage response funneling through TRP53 and TRP63. Some, but not all of the damage signaling to these proteins is transduced by CHK2, and we provide molecular evidence that CHK1 can also perform this function (see model in Fig. 3). The relative contributions of these transducer kinases in meiotic DNA damage responses is unclear, because the essential nature of CHK1 in embryonic and premeiotic germ cell development (Abe *et al.* 2018) complicates analyses. Nevertheless, CHK1 conditional mutagenesis and depletion experiments indicate that this kinase plays a role in modulating cell cycle progression in spermatocytes during meiotic prophase I (Abe *et al.* 2018), and in oocytes at the G2/M checkpoint (Chen *et al.* 2012). A key remaining question is whether CHK1 and CHK2 are the sole direct responders for TRP53 and TRP63, or if another transducer kinase(s), such as casein kinases 1 or 2 (CK1, CK2), function in parallel (Fig. 3).

**Figure 3.**
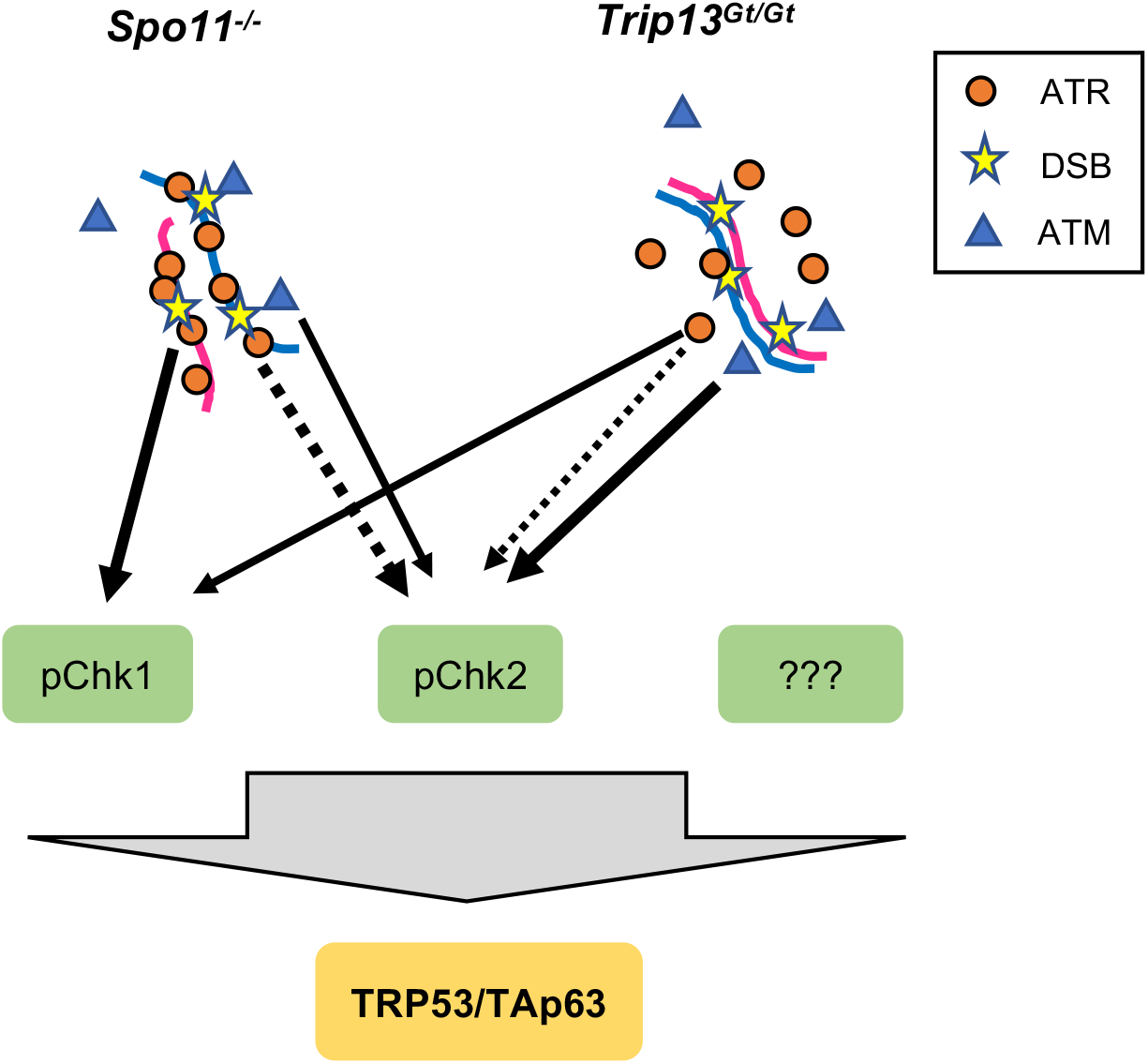
Model of checkpoint signaling in mouse oocytes. We propose that all DSB damage signaling in oocytes requires activation of TRP53 and TAp63 for complete oocyte elimination. The dashed lines represent non-canonical phosphorylation of CHK2 by ATR, and the thickness of all lines represents the relative amounts of activation in the two indicated mutant situations. We propose that in highly asynaptic *Spo11* mutant oocytes, the “pre-loading” of ATR as part of the MSUC response leads it to play a larger role in signaling to CHK1 and CHK2 than under situations in which DSBs occur on synapsed chromosomes.

## Acknowledgments

This work was supported by NIH grant GM45415 to J.C.S. and an institutional training grant (T32HD057854) that supported J.C.B. A. Mills originally provided us with TAp63 mutant mice.

## Materials and Methods

### Mice

Alleles used in this study and their genetic backgrounds were previously described ((Bolcun-Filas *et al.* 2014). Comparisons of compound mutants and controls utilized littermates whenever possible, otherwise animals from related parents or different litters from the same parents were used. Animal work was approved by Cornell’s Institutional Animal Care and Use Committee, under protocol 2004-0038 to JCS.

### Histology and Follicle Quantification

Ovaries were fixed in Bouin’s, embedded in paraffin, serially sectioned at 6μm, and stained with hematoxylin and eosin (H&E). Follicle identification (Myers *et al.* 2004) and quantification was as described (Bolcun-Filas *et al.* 2014). Graphs and statistical analysis were performed with GraphPad Prism8. Comparisons of follicle numbers across genotypes were performed using an Ordinary one-way ANOVA test.

### Western Blot Analysis of Protein Phosphorylation

Ovaries from postnatal (3-5 day old) mice were collected and divided into control and treatment groups. Treated groups were exposed to 3Gy of ionizing radiation as described above and proteins were extracted 3 hours post irradiation. Ovaries from all the females in the litter were dissected and individually frozen while genotyping was performed. Proteins from ovaries of selected genotypes were pooled into groups of 4 and extracted with lysis buffer containing: 20mM Tris-HCl (pH 7.4), 150 mM NaCl, 1mM EDTA, 1mM EGTA, 1% Triton X-100, protease Inhibitors (Complete Mini-Roche#11836153001), and phosphoprotease Inhibitors (PhosSTOP-Roche# 04906845001). Proteins were resolved on 4-20% gradient acrylamide gels (Biorad, cat. #4561093), transferred to PVDF transfer membranes (Millipore, cat. # IPVH00010) and blocked with 5% BSA or 5% nonfat milk according to the manufacturer datasheet for the corresponding antibody. Membranes were probed with rabbit anti-Phospho-Chk1 (Ser345) (1:750, Cell Signaling 133D3), rabbit anti-phospho Ser15-p53 (rodent-specific 1:750, Cell Signaling D4S1H), and rabbit anti-DDX4/MVH (1:750, Abcam 13840).

